# Comparison of spontaneous recurrent seizures in rats following status epilepticus induced by organophosphate paraoxon, DFP, and sarin

**DOI:** 10.1101/2023.05.10.540087

**Authors:** Robert E. Blair, Elisa Hawkins, R. Pinchbeck Lauren, Robert J. DeLorenzo, Laxmikant S. Deshpande

**Author notes:** **To whom correspondence should be addressed:** Laxmikant S. Deshpande, M.Pharm., Ph.D., FAES., Virginia Commonwealth University, School of Medicine PO Box 980599, Richmond, VA 23298.

## Abstract

Organophosphate (OP) compounds are highly toxic and include household, industrial, agricultural, and chemical warfare nerve agents (CWNA). OP exposure inhibits acetylcholinesterase enzyme, causing cholinergic overstimulation that can evolve into status epilepticus (SE) and produce lethality. Furthermore, OP-SE survival is associated with mood and memory dysfunction and spontaneous recurrent seizures (SRS). Here we assessed hippocampal pathology and chronic SRS following SE induced by OP agents in rats. Male Sprague-Dawley rats were injected with 1.5x LD_50_ of various OP agents, followed by atropine and 2-PAM. At 1-h post-OP-SE onset, midazolam was administered to control SE. Approximately 6 months following OP-SE, SRS were evaluated using continuous video-EEG monitoring. Histopathology was conducted using Hematoxylin and Eosin (H&E), while silver sulfide (Timm) staining was utilized to assess Mossy Fiber Sprouting (MFS). Over 60% of OP-SE surviving rats developed SRS with varying seizure frequencies, durations, and Racine severity scores. H&E staining revealed a significant hippocampal neuronal loss, while Timm staining revealed extensive MFS within the inner molecular region of the dentate gyrus of SRS-expressing OP-SE rats. This study demonstrates that OP-SE is associated with hippocampal neuronal loss, extensive MFS, and SRS, all hallmarks of chronic epilepsy.

## 1. Introduction

Organophosphate (OP) chemicals include household pesticides and herbicides, industrial solvents and chemical intermediates, and agricultural insecticides and pesticides (Costa, 2018). OP compounds also include chemical warfare nerve agents (CWNA), and civilian and military personnel have experienced mass OP exposure events (Newmark, 2004; Chao et al., 2010; BBCNews, 2013; Sugiyama et al., 2020). Unfortunately, accidental contamination of food products with OP agents leads to mass OP poisoning events, commonly reported in developing countries (Alvaro Javier, 2014; Mohiuddin et al., 2016). Further, OP pesticides represent a common source of poison for suicide attempts in developing nations (Karunarathne et al., 2021). Thus, OP exposure could occur under domestic, occupational, intentional, warfare, or terrorism-related scenarios.

Mechanistically, OPs are inhibitors of the cholinesterase (ChE) enzyme. In the brain, OP-induced inhibition of acetylcholinesterase (AChE) leads to rapid accumulation of acetylcholine (ACh) at the synapses, producing acute cholinergic symptoms such as headache, confusion, miosis, lacrimation, salivation, loss of sphincter tone, and muscle fasciculations (Bajgar, 2004; Tsai and Lein, 2021). Depending on the dose and OP type, these symptoms could rapidly evolve into unremitting seizures (status epilepticus, SE) followed by bradycardia, respiratory distress, and death if emergency intervention is not possible (Tattersall, 2009; Chuang et al., 2019). The current mainstay of OP poisoning treatment (Finkelstein et al., 1989; Eddleston et al., 2008; Chemical Hazards Emergency Medical Management, 2013) uses the 3-drug standard-of-care (SOC) approach consisting of 1) atropine, a muscarinic antagonist, to dry the copious secretions in the airway, 2) pralidoxime or 2-PAM, an AChE reactivator that would help catalysis of accumulated synaptic acetylcholine, and finally, 3) a benzodiazepine such as midazolam to control the SE.

Despite these advances, the effectiveness of current treatments is limited to the acute care setting (Jett and Spriggs, 2020; Jett and Laney, 2021). When the first responders arrive at the scene in mass civilian OP exposures, the immediate goal is to treat the cholinergic symptoms and control SE. SE is a clinical emergency associated with high mortality and morbidity (Towne et al., 1994; Shorvon, 2013). SE is known to produce multifocal neuronal injury (Sankar et al., 1998) and is also a significant risk factor for developing recurrent seizures or acquired epilepsy (Hesdorffer et al., 1998). Long-term SE and epilepsy outcomes include neurological morbidities, depression (Fiest et al., 2013), and cognitive deficits (Helmstaedter, 2007). Experimental models to study chemical-induced SE outcomes have been developed in laboratory settings using both OP (Deshpande et al., 2010; Deshpande et al., 2014a; Guignet et al., 2020; Reddy et al., 2021; Rojas et al., 2021) and non-OP chemoconvulsants (Covolan and Mello, 2000; Leite et al., 2002; Groticke et al., 2008; Mazarati et al., 2008). Considering the effectiveness of the current treatment, it is hopeful that a significant proportion of exposed individuals will survive OP exposure; however, many OP-SE survivors will inevitably develop chronic neurological morbidities.

Investigators have utilized various OPs, including pesticide surrogates such as paraoxon (POX) (Deshpande et al., 2014a; Deshpande et al., 2014b), CWNA surrogates such as diisopropyl fluorophosphate (DFP) (Deshpande et al., 2010; Liu et al., 2012; Flannery et al., 2016; Pouliot et al., 2016; Wu et al., 2018; Guignet et al., 2020; Calsbeek et al., 2021; Rojas et al., 2022), and CWNAs themselves, like sarin (GB, O-isopropyl methylphosphonofluoridate) (Shih et al., 2003; Chapman et al., 2006; Damodaran et al., 2006; Grauer et al., 2008) and soman (GD) (Myhrer et al., 2005; Angoa-Perez et al., 2010; de Araujo Furtado et al., 2012; Perkins et al., 2016; Gage et al., 2021), to study OP exposure’s acute and chronic effects. These models report neuronal injury and neuroinflammation in multiple brain regions and are associated with SRS and behavioral morbidities such as mood and memory impairments (de Araujo Furtado et al., 2010; Shrot et al., 2014; Bar-Klein et al., 2016; Gage et al., 2020; Guignet et al., 2020; McCarren et al., 2020). However, these experimental studies did not always use a lethal intoxicating OP dose or implemented the 3-drug SOC countermeasures at human-equivalent doses. Additionally, previous studies did not always implement benzodiazepine “rescue” to control OP-SE, allowing for the assessment of both sub-acute seizure activity (days-weeks) and the long-term emergence of SRS following OP-SE. In this study, we incorporated these modeling variables, and for the first time report the occurrence of MFS in association with hilar neuronal loss and chronic SRS following OP-SE.

## 2. Methods

### 2.1 Drugs and Chemicals

Paraoxon (POX), Atropine sulfate, and Pralidoxime chloride (2-PAM) were obtained from Millipore-Sigma (St. Louis, MO). DFP was obtained from Chem Service Inc (West Chester, PA). Midazolam was obtained from VCU Health System Pharmacy. All the drugs were prepared fresh on the day of the experiment.

### 2.2 Animals

All animal use procedures were in strict accordance with the National Institute of Health Guide for the Care and Use of Laboratory Animals and approved by Virginia Commonwealth University’s Institutional Animal Care and Use Committee (IACUC). In addition, GB studies conducted at MRI Global (Kansas, MO) were reviewed and received appropriate approval from their IACUC. Male Sprague-Dawley rats were obtained from Envigo (Indianapolis, IN) at 9 weeks of age. Animals were housed two per cage at 20-22°C with a 12 Light: 12 Dark hour cycle (lights on 0600-01800 h) and free access to food and water.

### 2.3 Electrode implantation

The EEG-video acquisition was utilized to analyze both behavioral and electrographic seizures. At least 6 weeks following OP-induced SE, animals underwent surgical implantation of skull surface electrodes. Under general anesthesia with isoflurane/O2 (5% induction; 2.5% maintenance), bupivacaine (0.5%, 0.1 ccs, s.c.) was administered along the incision site for local anesthesia, and a 15 mm sagittal incision of the scalp was made along the midline to expose the skull. Rats were stereotaxically implanted with four electrode screws attached to Teflon-insulated stainless steel MedWire® (Plastics One, Roanoke, VA, USA.). Electrode screws were positioned through burr holes to contact but did not penetrate the dura mater above the right and left frontal and motor cortices (AP, ±3 mm and ML, ±3 mm from bregma); a fifth electrode screw was positioned over the cerebellum to serve as a reference, and two additional (non-electrode) skull screws were inserted for structural support. Amphenol terminal pins for each electrode were seated into a threaded electrode pedestal (Plastics One, Roanoke, VA). This assembly was secured to the skull with dental acrylate, and the incision site was closed with surgical adhesive. Rats were administered meloxicam SR (4 mg/kg, s.c.) to provide 72 hours of analgesia and were allowed 2 weeks of recovery time before the start of EEG-video recordings.

### 2.4 Induction of SE with POX, DFP, and GB

Separate cohorts of rats were injected with either POX (2 mg/kg, s.c.) (Deshpande et al., 2014a) or DFP (4 mg/kg, s.c.) (Deshpande et al., 2010). One minute later, immediate prophylactic treatment with atropine (0.5 mg/kg, i.m.) and 2-PAM (25 mg/kg, i.m.) was instituted. OP exposure produced signs of progressive cholinergic hyperstimulation culminating in continuous seizures. A Racine score of 4-5 (rats rearing and falling on their backs and seizing uncontrollably) was the criteria to mark the SE onset 5-10 mins post OP injection. At 1-h post-SE onset, midazolam (1.78 mg/kg, i.m.) was administered to control SE. These rat doses were selected using human equivalents calculated using a human-to-rat dose translation equation(Reagan-Shaw et al., 2008; Nair and Jacob, 2016). Experiments with sarin (GB) were conducted at MRI Global facility (Kansas, MO). GB was dissolved in ice-cold saline and injected subcutaneously at 132 μg/kg. One minute later, rats were injected with atropine methyl nitrate (2 mg/kg, i.m.) and 2-PAM (25 mg/kg, i.m.). Rats displayed severe hypercholinergic symptoms, including Racine 4-5 seizures. SE rats were treated with midazolam (5 mg/kg, i.m.) at one and three hours following the SE onset. All OP-SE rats received supportive care, including saline injection, oral glucose/ milk supplementation, and liquid chow (diet gel packs) in their cages to aid recovery until they increased weight. MRI Global shipped GB rats along with age-matched controls to VCU one week following the termination of SE. All rats were housed one/ cage with appropriate environmental enrichment following OP-SE inside the VCU vivarium.

### 2.5 Video and EEG-video evaluation of epileptic seizures

For EEG-video acquisition, 6-channel wire leads were securely connected to the threaded electrode pedestal on the rat and then connected to an electrical-swivel commutator (Plastics One, Roanoke, VA) to allow for unrestricted movement of the animal while maintaining the continuity of biopotential signals. EEG signals were amplified using a Grass model 8–10D (Grass Technologies, West Warwick, RI) and digitized using a Powerlab 16/30 data acquisition system (AD Instruments, Colorado Springs, CO). Acquisition and offline evaluation of digitally acquired EEG were carried out with Labchart-7 software (AD Instruments, Colorado Springs, CO). Synchronized with EEG, a time-stamped video was obtained using a high-resolution CCD camera (C20-DW-6; Pelco, Fresno, CA) with a wide dynamic range allowing for imaging at both daylight and dark cycle under infrared illumination. Digital video was acquired and reviewed offline using a digital video capture card and software (GeoVision Inc., Irvine, CA). Animals underwent continuous EEG-video monitoring for at least 7 days and were allowed to move freely within the cage with ad-lib access to food and water. Electrographic seizures are defined by discrete epileptiform events characterized by episodes of high frequency (>2/sec) and increased voltage multi-spike complexes and/or synchronized spike or wave activity lasting >10sec. We do not classify interictal spikes or epileptiform discharges that last less than 10 sec as seizures. Duration of electrographic seizures was analyzed offline and measured from the onset of low amplitude high-frequency spike activity, which transitioned into high amplitude multi-spike burst activity. The termination of a seizure was indicated by both a return of the EEG to inter-seizure baseline activity and cessation of video-behavioral convulsant activity. Two video-only sessions of control and OP-treated animals were recorded for 3–5 days at a minimum of 2 months apart, and behavioral seizures were evaluated offline. For video-only and EEG-video seizure analysis, the severity of each behavioral seizure was scored on a scale of 0-5, according to Racine (Racine, 1972). The criteria for the behavioral seize score were: 0 - behavioral arrest, hair raising, excitement, and rapid respiration; 1 - mouth movement of lips/tongue, vibrissae movement, and salivation; 2 – head “bobbing”/clonus; 3 – forelimb clonus; 4 – clonic rearing, and 5 – clonic rearing with loss of postural control. All electrographic seizures detected had a behavioral seizure score of ≥ 2.

### 2.6 Histology

Control and OP-treated rats underwent perfusion/fixation four to six months after OP-induced SE. Following induction of deep anesthesia with ketamine/xylazine (75 mg/ 7.5 mg/kg, i.p.), rats were perfused transcardially with isotonic saline followed by 100 ml of 1.2% NaS and then perfused with 250 ml of 4% paraformaldehyde in a 0.1 M phosphate buffer (pH 7.4). Brains were allowed further fixation for 24 h in buffered 4% paraformaldehyde at 4°C and then cryoprotected in a 30% sucrose in 0.1 M phosphate buffer (pH 7.4) for 3 days at 4°C. Brains were then embedded in OCT by snap freezing in isopentane (−15° C). Coronal sections (20 microns) were prepared using a Leica CM3050S cryostat (Leica Biosystems, Deer Park, IL) and adhered to glass slides (Superfrost Plus; Fisher Scientific, Pittsburg, PA) and stored at -80C until use. Following staining with either Hematoxylin and eosin (H&E) or the Timm method, stained sections were visualized using an Olympus IX70 microscope under brightfield illumination with 10X and 20X objectives. Digital greyscale images of stained sections were acquired with a CCD camera (ORCA-ER; Hamamatsu Corp, Bridgewater, NJ) using MetaMorph image analysis software (Molecular Devices, San Jose, CA). All settings for illumination, image acquisition, and analysis were consistent throughout each stained section evaluated.

### 2.7 Mossy Fiber Sprouting

Mossy Fiber Sprouting (MFS) was evaluated using the Timm method, which labels mossy fiber synaptic terminals which are high in Zinc (Frederickson and Danscher, 1990). Sections containing dorsal hippocampus from both control and OP-treated rats were equilibrated to room temperature and then placed in ddH2O for 5 minutes, followed by incubation in the developer containing: 150 ml 50% gum Arabic, 25 ml of citrate buffer (6.4 g citric acid, 5.9 g sodium citrate q.s. to 25 ml in ddH2O), 75 ml of hydroquinone (4.2 g q.s. to 75ml in ddH2O), and 1.25 ml of a 15% silver nitrate. Staining was developed in the dark for 60-90 minutes. Stained sections were washed twice in ddH2O, underwent alcohol dehydration, and cleared in Xylenes and coverslip with permount. The intensity of Timm-stained granules within the inner molecular layer (IML) of the DG dorsal blade between the tip and the crest from the right and left hemispheres were scored on a scale of 0-5 as previously described (Sutula et al., 1996) employing the following criteria: 0– no granules in the IML, 1– sparse granules in the IML with non-continuous distribution, 2– increased granule density in a continuous pattern throughout the IML, 3– increased granule density in a continuous pattern with patches of confluent granules, 4– a dense and confluent band of granules throughout the IML, and 5-a prominent and dense band of confluent granules that innervate into the IML. Two blinded observers scored MFS severity.

### 2.8 Neuronal Cell Counts

Hematoxylin and eosin (H&E) staining of sections was used to evaluate neuronal loss within the hilar region of the DG using Mayer’s H medium and alcoholic eosin Y (Sigma-Aldrich, Saint Louis, MO). Briefly, air-dried sections were equilibrated for 60 sec in ddH2O, followed by incubation in Mayer’s H medium for 5 min. Sections were then washed in ddH2O, followed by bluing in Scott’s tap water for 5 min. Following a brief wash, sections were stained in alcoholic E medium for 60 sec, dipped five times each in 50% and 70% EtOH sequentially, dehydrated, cleared in xylenes and coverslip with permount.

Grayscale 16-bit images of H&E stains were analyzed using ImageJ software (NIH – public domain) to determine neuronal cell counts within the inner hilar region of the DG. The inner hilar region analyzed was determined by bisecting between the inner infrapyramidal and suprapyramidal granule cell layers just medial to the termination of the CA3 region. It included the hilar region to the septal apex of the DG while excluding the subgranular zone. The mean grey background level was measured within the DG hilus in a region that excluded any cell/punctate staining and set as the maximum greyscale level in the B&C tool, while the minimum level was set to a constant value of 1260 for all images analyzed. These levels were applied to remove the background, binarized the image, and then auto-thresholded. Cells within the threshold were measured using the analyze particle module with a set pixel size range to detect only stained hilar neurons. Cell counts were represented as cells/mm^2^ within the inner hilar region measured. Hilar cell counts of the left and right hemispheres were made from three slides/sections per animal, including the anterior, medial, and posterior dorsal hippocampus.

### 2.9 Data analysis

Data analysis was accomplished using SigmaPlot 14.0 software. Data represented as mean ± SEM. A student’s t-test was employed to compare respective age-matched controls and POX, DFP, or GB rats. A p-value of 0.05 was considered to note statistical significance.

## 3. Results

### 3.1 Chronic SRS following OP-induced SE

At 6 months post OP-SE, video-EEG monitoring revealed the presence of SRS in surviving rats. A typical SRS recorded on EEG consisted of seizure onset indicated by rapid spike discharges with frequencies between 10-20 Hz, followed by well-formed polyspikes and slow-wave discharges, corresponding to the tonic and clonic phases of the seizure activity, respectively. The behavioral correlates of these seizures consisted of a sudden cessation of activity associated with vacuous chewing, facial twitching, head jerks, and forelimb clonus. Most observed seizures were generalized convulsive.

In the POX group, video-EEG monitoring revealed the presence of SRS in 65% of POX-SE rats (n= 19 out of 29). The average seizure duration was 48.7 ± 3.7 s. POX rats exhibited seizure frequency at 8.6 ± 1.9 seizures/day with a severity score of 3.79 ± 0.18, as noted using the Racine scale. In the DFP group, video-EEG monitoring revealed the presence of SRS in 67% of DFP-SE rats (n= 18 out of 27). DFP rats exhibited a seizure frequency of 15.62 ± 2.6 seizures/day. The average seizure duration was 60.83 ± 12.8 s. The Racine seizure severity was 3.38 ± 0.14. In the GB (sarin) group, video-EEG monitoring revealed the presence of SRS in 61% of GB-SE rats (n= 17 out of 28). GB rats exhibited a seizure frequency of 10.99 ± 2.5 seizures/day. The average seizure duration was 44.97 ± 4.5 s, while the seizure severity was 3.29 ± 0.28.

### 3.2 Neuronal cell loss following OP-induced SE

H&E stain of age-matched control animals showed the presence of pyramidal-shaped neurons throughout the DG hilus region. At the same time, there was a significant decrease in neuronal cell number within the hilus from animals that developed SRSs following exposure to POX, DFP, and Sarin (Fig. 3). analysis of hilar cell counts/mm^2^ in control were (102.8 ± 9.0), while a significant decrease following OP-induced SE with development of SRSs was observed for POX (69.3 ± 11.4), DFP (62.5 ± 11.7), and GB (51.2 ± 6.8) (*p<0.05, Student’s t-test, n= 4-6 rats/ group).

**Figure 1.**
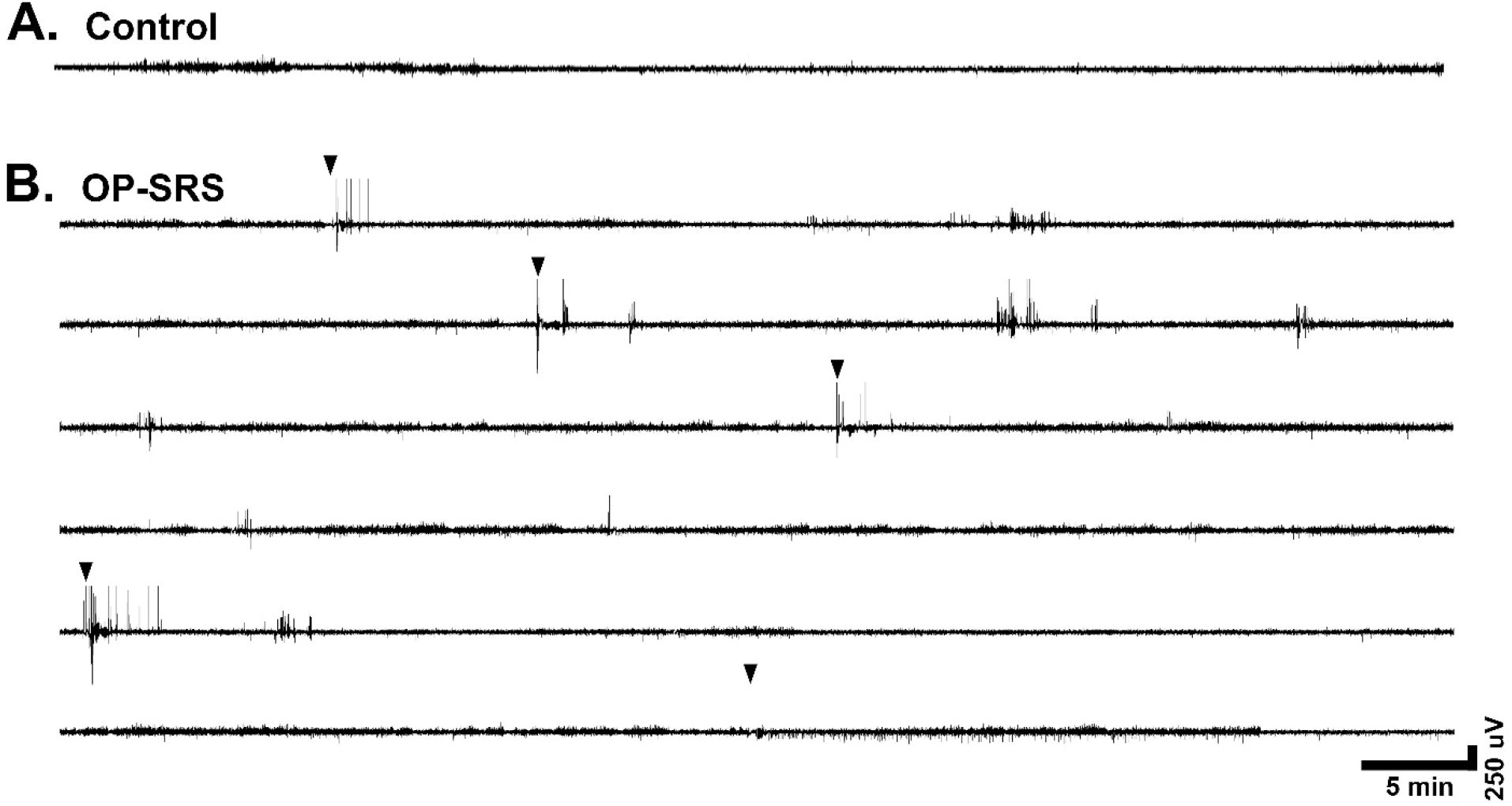
Development of long-term SRS in rats following survival from OP-SE. **A**. Representative 60 min recording of EEG from an age-matched control rat. **B**. Representative 6 h recording from an SRS expressing rat previously exposed to OP-SE (POX). This rat displayed SRS activity as indicated by 5 epileptic seizures (arrowheads) during a 6 h recording. DFP and GB rats also manifested SRS with variations in SRS duration and frequencies.

**Figure 2.**
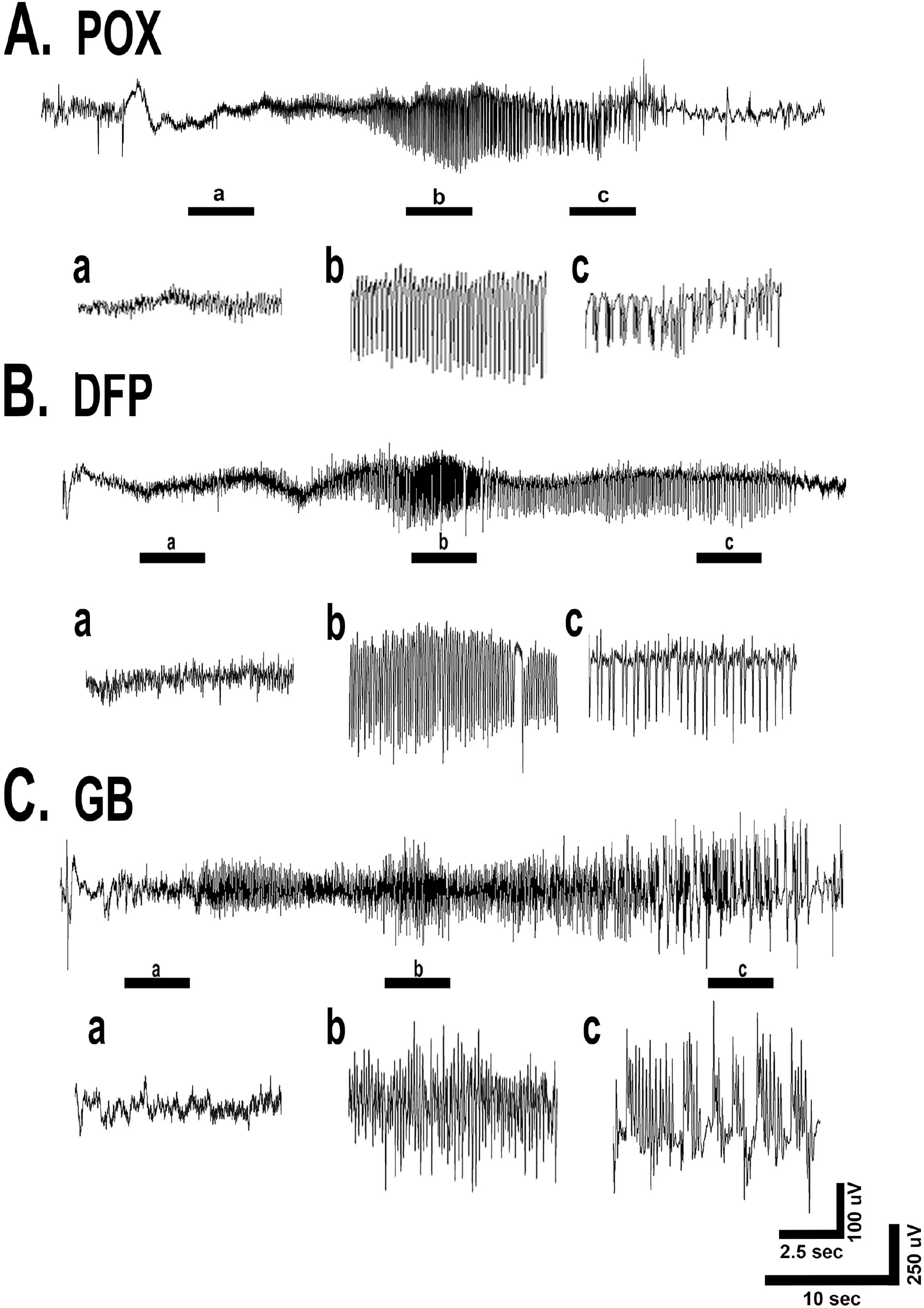
Chronic SRS around 4-6 months after OP-SE. EEG recording from hippocampal surface electrodes from OP rats manifested SRS activity. **A**. EEG recording from a POX-SE rat with three segments expanded to indicate various SRS characteristics. These indicate **a**. high-frequency low-voltage seizure onset, **b**. high-voltage, high-frequency tonic spike discharges, followed by **c**. well-formed polyspike and slow wave discharges, correspond to the seizure activity’s tonic-clonic phase. **B**. EEG recording from a DFP rat. **C**. EEG recording from a GB rat. Similar to the SRS observed in the POX rats, DFP and GB rats also manifested SRS with similar EEG characteristics.

**Figure 3.**
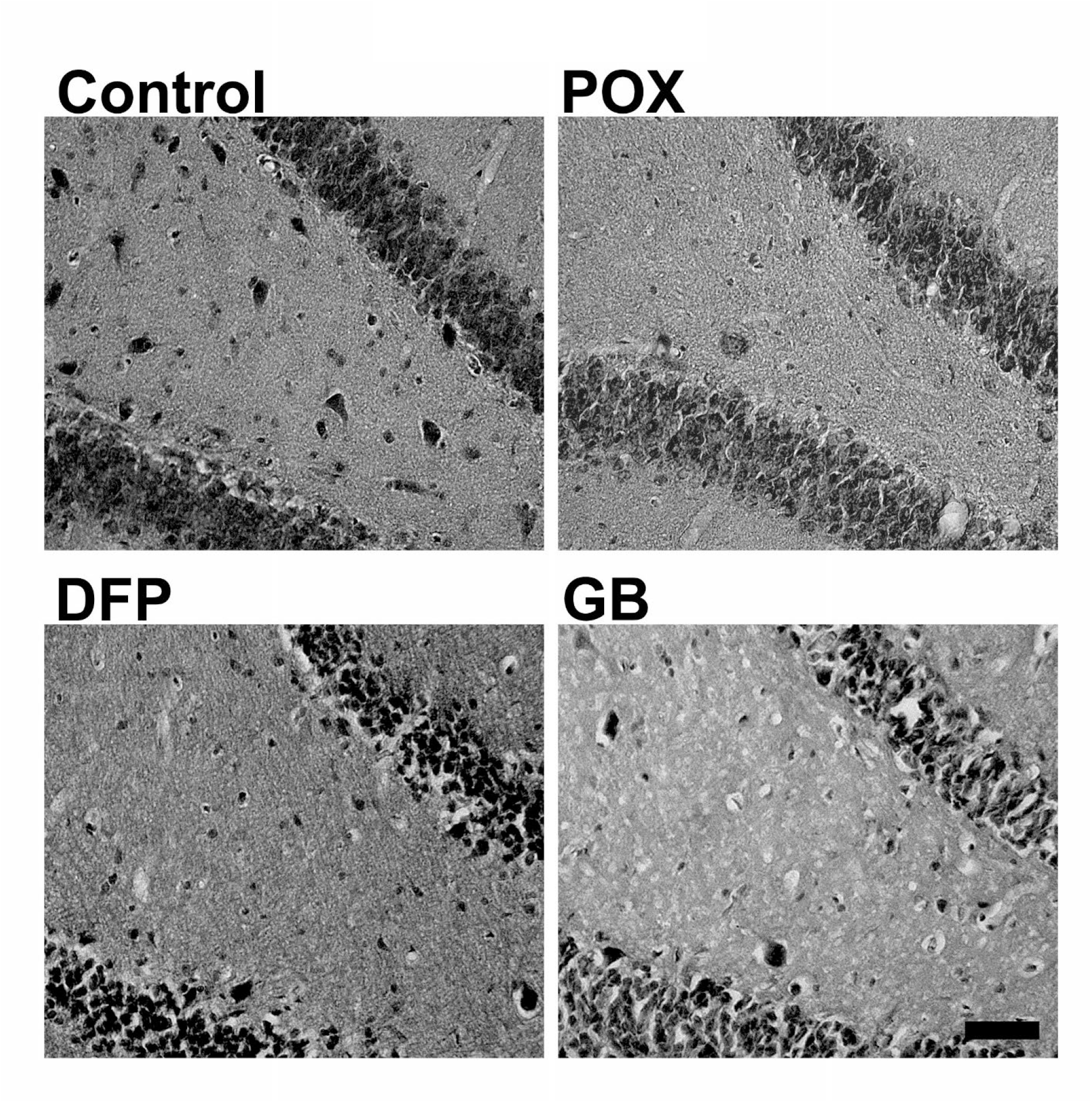
Hippocampal neuronal loss in SRS expressing OP-SE rats. H & E-stained brain sections of rats 6-m after OP exposure. Representative staining from an age-matched control rat showed well-defined staining of multiple pyramidal-shaped neurons within the inner hilar region of the DG. In contrast, a significant loss of hilar neuronal cells in the DG from POX, DFP, and GB-SE animals was observed. Scale = 100 microns

### 3.3 Mossy Fiber Sprouting Following OP-induced SE

Age-matched control animals show no or barely detectable Timm-stained granules of mossy fiber synaptic terminals within the inner molecular region of the DG. Timm staining revealed the presence of a dense, continuous band of MFS within the IML of the dentate gyrus of SRS-expressing OP rats but not control rats. Exposure to all three OPs (POX, DFP, and sarin) with subsequent development of SRSs showed MFS (fig. 4). Mean Timm scores were significantly higher in rats following exposure to POX (2.75 ± 0.704), DFP (4.5 ± 0.5), and GB (3.0 ± 0.87) when compared to control rats (0.1 ± 0.083) (*p<0.05, Student’s t-test, n= 3-6 rats/ group).

**Figure 4.**
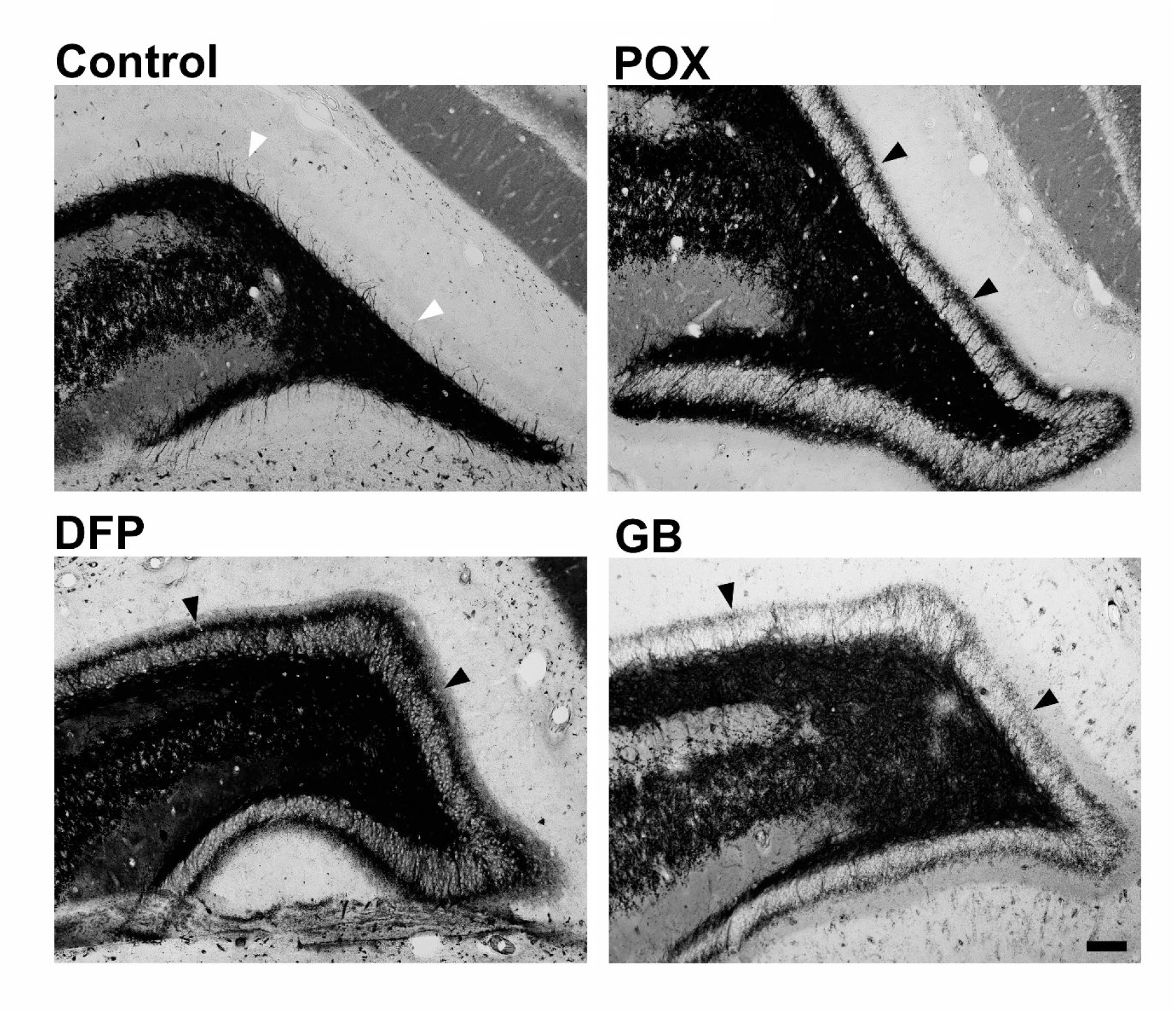
Mossy fiber sprouting in SRS expressing OP-SE rats. Representative Timm stains around 6-m after OP-SE revealed the presence of a dense, continuous band of MFS (black arrowheads) originating from the granule cells and innervating the IML of the dentate gyrus of POX, DFP, and GB rats with SRS. Timm stain from age-matched control animals reveals little to no MFS within the IML of the DG (white arrowheads). scale = 100 microns

## 4. Discussion

Our studies demonstrated that SE following exposure to lethal concentrations of OP agents leads to chronic SRS, significant neuronal loss, and extensive MFS in the hilus and the IML of the dentate gyrus, respectively, all hallmarks of chronic epilepsy. OP agents are considered threat agents and have been weaponized previously and used with lethal outcomes against the military and civilian populations. The availability of FDA-approved SOC treatments has significantly improved mortality, yet, long-term neurological morbidities remain a significant concern in OP-SE survivors. Using three different OP agents, which include a pesticide surrogate (POX), an OP threat agent (DFP), and a CWNA (GB), we provide a model system for studying chronic aftermaths of OP toxicities.

We standardized the model using the following criteria for all three OPs. 1) We utilized 1.5x LD_50_ concentration to simulate lethal intoxication. 2) We provided emergency treatment using all three FDA-approved standards of care, namely, atropine, 2-PAM, and midazolam. 3) The SOC treatment utilized human-equivalent doses. 4) We controlled seizures at 1-h post-SE-onset to simulate a delay in emergency care during a mass exposure scenario. 5) We evaluated electrographic and behavioral SRS, conducted histopathology, and assessed MFS at 6-m after the termination of OP-SE to truly understand long-term, chronic effects of OP-SE on seizure outcomes. Using this paradigm, our data revealed that over 60% of OP-SE rats develop chronic SRS, exhibit significant hilar neuronal loss, and display profound MFS, all indicative of temporal-lobe epilepsy-like pathology in OP-SE surviving rats.

The development of SRS or chronic acquired epilepsy following survival from OP-SE has been previously reported. For example, a pair of studies showed that adult male Sprague-Dawley rats poisoned with POX (450 μg/kg, i.m.) and immediately treated with atropine (3 mg/kg, i.m.) and obidoxime (20 mg/kg, i.m.) and 30-mins later with midazolam (1 mg/kg, i.m.) resulted in around 50-55% of rats developing SRS at 4-7 weeks post-SE (Shrot et al., 2014; Bar-Klein et al., 2016). In another study, adult male Sprague-Dawley rats were initially given a reversible AChE inhibitor pyridostigmine bromide (0.1 mg/kg, i.m.) 30 min before administration of DFP (4 mg/kg, s.c.) which was immediately followed by atropine sulfate (2 mg/kg, i.m.) and pralidoxime (25 mg/kg, i.m.). No midazolam intervention occurred in this study, where SE lasted several hours. Around 70% of rats exhibited SRS over the first 2 months post-exposure (Guignet et al., 2020). Yet another study looked at SRS in male and female rats four weeks post-DFP exposure. Here DFP (3-4 mg/kg, s.c.) was injected, followed by atropine sulfate (2 mg/kg, i.m.) and 2-PAM (25 mg/kg, i.m.). Animals displayed SE for two hours before diazepam (5 mg/kg, i.m.) was administered to control SE. Around 50% of male and 30% of female rats showed SRS in the four weeks post-DFP exposure (Gage et al., 2020). CWNA agent soman (GA) has also been studied for the emergence of SRS following SE. The mouse GA model was associated with high SE mortality (around 40%) and displayed SRS in 100% of subjects assessed six weeks post-SE (McCarren et al., 2020). The rat GA model was associated with lower acute mortality but also exhibited a lower SRS development rate of around 29% from 5-10 days following SE (de Araujo Furtado et al., 2010). Our studies support these observations and provide additional data on SRS characteristics at long-term time points post OP-SE and robust validation using standardized experimental modalities.

MFS (Franck et al., 1995; Proper et al., 2000; Buckmaster et al., 2002; Buckmaster, 2014) and cell loss in the hilus region of the dentate gyrus (Covolan and Mello, 2000; Rojas et al., 2021; DeFelipe et al., 2022) is reported in both human epilepsy and animal models of epilepsy. Following SE, hippocampal mossy fibers of dentate granule cells are said to develop recurrent collaterals that invade the dentate IML to generate recurrent excitation by forming granule cell-granule cell synapses (Mello et al., 1993; Okazaki et al., 1995). Our studies indicated profound MFS in all OP-SE rats that exhibited SRS. These maladaptive mossy fiber collaterals originate from the dentate granule cells, which cross through and innervate their own dendritic fields within the IML. This MFS creates an aberrant, recurrent, excitatory circuit that is thought to underlie SRS following SE (Buckmaster, 2014). While we did not find a correlation between the extent and severity of MFS and SRS frequency, MFS was noted only in the animals that developed SRS following OP-SE.

Interestingly, hilar cell loss was noted in all the rats that experienced OP-SE, irrespective of SRS outcomes. This suggests that MFS could be one of the factors for developing chronic epilepsy following OP-SE. Hilar cell loss and MFS represent hippocampal structural changes that possibly underlie SRS expression following OP-SE. Identifying the circuit mechanisms for SRS is critical for understanding the chronic pathology and development of pathological synaptic plasticity and long-term neurological morbidities following survival from OP SE.

In this study, we focused on assessing the chronic nature of SRS in OP-SE surviving rats. Chronic SE morbidities, surgical complications, and dislodging of headsets significantly contribute to animal attrition, especially when long-term studies are planned. We implanted electrodes and monitored for EEG 6 months post-OP-SE to conserve animal numbers. Thus, we did not study the emergence and progression of SRS in the period immediately following OP-SE. Furthermore, only male rats were utilized for OP-SE. Sex differences have been reported for OP toxicities. Thus, additional studies are needed to determine how female rats respond to OP-SE with subsequent SRS and hippocampal pathology.

In conclusion, this study demonstrates that SE induced by various OP agents leads to neuron loss, MFS, and SRS development, which are hallmarks of epilepsy. These rat OP SE-induced SRS models offer a valuable resource to evaluate the effects of lethal OP intoxication on the development of chronic epilepsy and behavioral comorbidities. These models are also helpful in studying molecular mechanisms underlying the development of OP comorbidities and identifying novel treatments for OP toxicities.

## Acknowledgments

We thank the members of the MRI Global team with GB exposures. This work was supported by the Countermeasures Against Chemical Threats (CounterACT) Program, the National Institutes of Health (NIH) Office of the Director (OD), and the National Institute of Neurological Disorders and Stroke (NINDS) [Grant U01NS1058]. Its contents are solely the responsibility of the authors and do not necessarily represent the official views of the federal government.

